# Engineered model of t(7;12)(q36;p13) AML recapitulates patient-specific features and gene expression profiles

**DOI:** 10.1101/2022.06.14.496084

**Authors:** Denise Ragusa, Ylenia Cicirò, Concetta Federico, Salvatore Saccone, Francesca Bruno, Reza Saeedi, Cristina Sisu, Cristina Pina, Arturo Sala, Sabrina Tosi

## Abstract

Acute myeloid leukaemia carrying the translocation t(7;12)(q36;p13) is an adverse-risk leukaemia uniquely observed in infants. Despite constituting up to 30% of cases in under 2-year-olds, it remains poorly understood. Known molecular features are ectopic overexpression of the *MNX1* gene and generation of a fusion transcript in 50% of patients. Lack of research models has hindered understanding of t(7;12) biology, which has historically focused on *MNX1* overexpression rather than the cytogenetic entity itself. Here, we employed CRISPR/Cas9 to generate t(7;12) in the human K562 cell line, and in healthy CD34+ haematopoietic progenitors where the translocation was not sustained in long-term cultures or through serial replating. In contrast, in K562 cells, t(7;12) was propagated in self-renewing clonogenic assays, with sustained myeloid bias in colony formation and baseline depletion of erythroid signatures. Nuclear localisation analysis revealed repositioning of the translocated *MNX1* locus to the interior of t(7;12)-harbouring K562 nuclei - a known phenomenon in t(7;12) patients which associates with ectopic overexpression of *MNX1*. Crucially, the K562-t(7;12) model successfully recapitulated the transcriptional landscape of t(7;12) patient leukaemia. In summary, we engineered a clinically-relevant model of t(7;12) acute myeloid leukaemia with the potential to unravel targetable molecular mechanisms of disease.

## Introduction

The t(7;12)(q36;p13) translocation is a recurrent chromosomal rearrangement uniquely associated with infant acute myeloid leukaemia (AML) (1,2). T(7;12) is the second most common cytogenetic abnormality in AML patients below the age of 2 years (3), and associates with a dismal prognosis. Structurally, t(7;12) is a balanced translocation disrupting the long arm of chromosome 7 (7q36.3) and the short arm of chromosome 12 (12p13.2), producing two derivative chromosomes, der(7) and der(12). The 7q36.3 breakpoint lies proximal to the *MNX1* gene, which is entirely relocated to der(12). The breakpoint on 12p13, on the contrary, disrupts the *ETV6* gene on its 5’ portion (4-6).

Mechanisms of leukaemogenesis associated with t(7;12) remain poorly understood. A *MNX1/ETV6* fusion mRNA transcript is produced from der(12) in approximately half of patients with t(7;12) (7-9), but a translated MNX1/ETV6 protein has never been detected. Introduction of the chimaeric transcript did not transform mouse bone marrow cells, suggesting that it may not contribute to the leukaemic phenotype (10).

A common feature among all t(7;12) patients is overexpression of the *MNX1* gene (7,9,11), which is proposed to result from disruption of its genomic positioning (7). The majority of reports on t(7;12)-mediated leukaemogenesis have focused on *MNX1* overexpression (10,12,13). In vitro, overexpression of *MNX1* in mouse haematopoietic stem cells (HSCs) did not result in expansion or increased survival capacity (10), and was shown to induce senescence (13). In human cord blood CD34+ haematopoietic stem and progenitor cells (HSPC), *MNX1* overexpression induced erythroid transcriptional programmes and reduced colony-forming capacity. *In vivo*, transplantations of *MNX1* overexpressing bone marrow cells did not cause leukaemia in the recipients, but resulted in an accumulation of *MNX1*-overexpressing cells in the megakaryocytic-erythrocyte fraction, but not in the granulocytic-monocyte nor in the mature lymphoid compartments (13).

Genome editing technologies can be harnessed for generation of chromosomal abnormalities (14-16). The CRISPR/Cas9 gene editing tool employs the endonuclease activity of the Cas9 enzyme to induce DNA double stranded breaks (DSB) at specific loci directed by guide RNAs (gRNAs). The induction of two simultaneous DSBs can lead to erroneous break repair and rejoining of the ‘wrong’ DNA ends, thereby forming a chromosomal translocation (17,18). Here, we sought to recreate the t(7;12) rearrangement in vitro using CRISPR/Cas9 in the attempt to uncover biological cues to its mechanisms in AML.

## Results and Discussion

By delivery of CRISPR/Cas9 ribonucleoprotein (RNP) complexes targeting clinically accurate breakpoints described by Tosi *et al*. (5) and Simmons *et al*. (4) (**Figure 1A; Supplementary Table 1**), we were able to achieve the t(7;12) rearrangement in the leukaemia cell line K562, as confirmed by fluorescence in situ hybridisation (FISH) (**Figure 1B-C; Supplementary Figure 2A-C**) and amplification of genomic fusion junctions by polymerase chain reaction (PCR) (**Figure 1D; Supplementary Figure 1**). While *MNX1* is already expressed in K562 cells (19), the translocation further upregulated its expression (**Figure 1E**), consistent with ectopic *MNX1* overexpression in patients (7,9,11). Altered nuclear positioning has been proposed as a significant mechanism of *MNX1* overexpression in t(7;12) leukaemia, with relocalisation of the der(12) containing the translocated *MNX1* gene to a more internal transcriptionally active position within the nucleus, where expression of *MNX1* can be triggered (7). We used nuclear localisation analysis to understand whether this phenomenon was recapitulated in the K562 model. FISH using a custom 3-colour probe allowed to distinguish both derivatives in interphase nuclei (**Figure 1F; Supplementary Figure 2D-F**) in order to calculate radial nuclear location (RNL) values for each locus with respect to their positioning towards the interior or peripheral nuclear space (**Figure 1F-G**). As described in patients, the introduction of t(7;12) in K562 cells resulted in the altered positioning of the der(12) towards the nuclear interior, consistent with the relative gain in *MNX1* expression (**Figure 1B**). Conversely, the der(7) containing the 5’ of *ETV6* was repositioned towards the periphery (**Figure 1G**), also in agreement with the patient analysis by Ballabio *et al*. (7). Nuclear organisation as a mechanism of regulation of *MNX1* expression was also identified in leukaemias harbouring different interstitial deletions of chromosome 7q (20). Specifically, the localisation of *MNX1* was shown to be dependent on the GC-content of the chromosomal band harbouring the proximal breakpoint, with *MNX1* being expressed when breakpoints affected GC-rich bands, which resulted in relocation of *MNX1* towards the nuclear interior; in contrast, GC-poor breakpoints resulted in localisation at the periphery and *MNX1* was not expressed (20).

**Figure 1.**
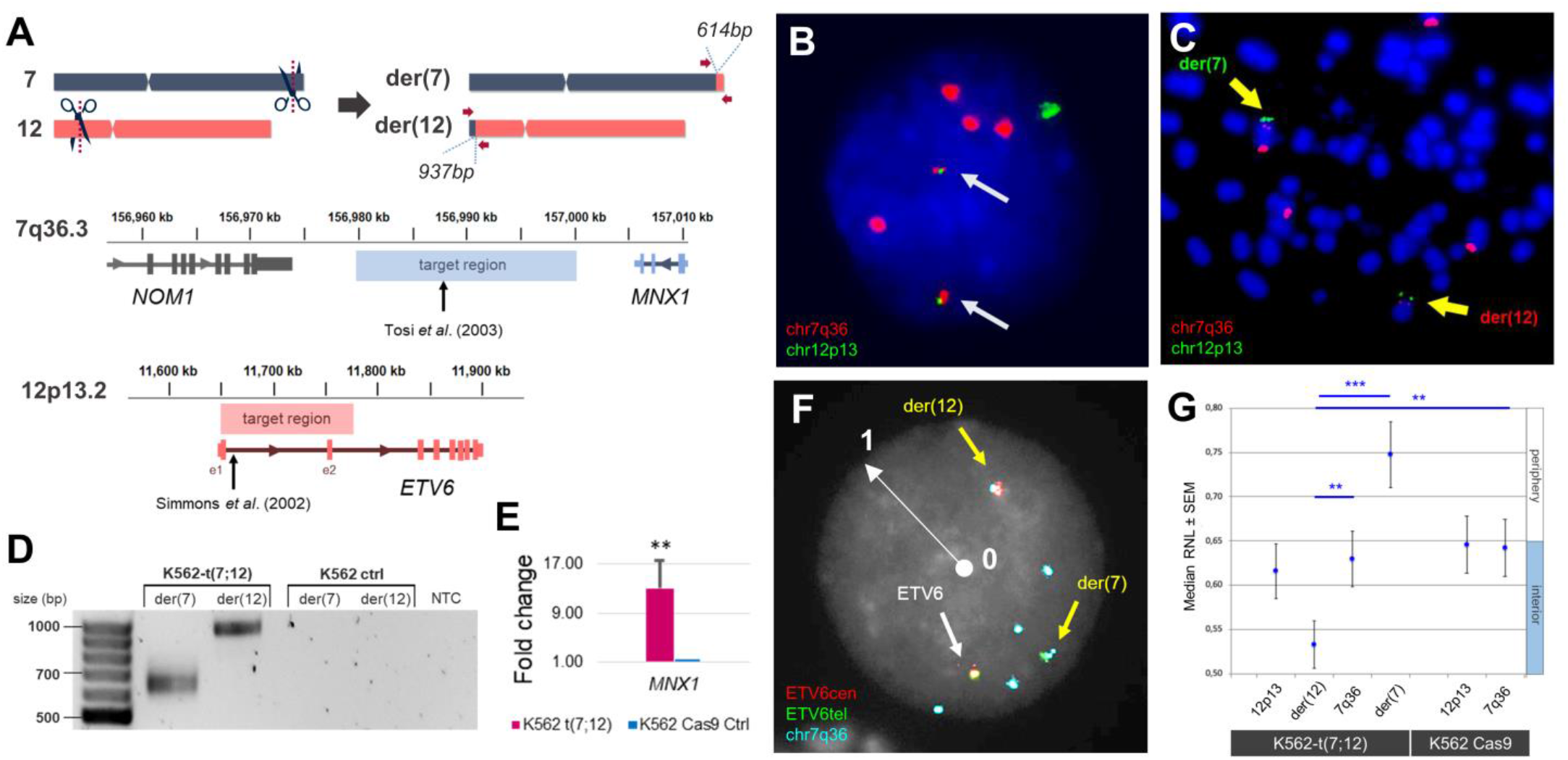
Generation of t(7;12)(q36;p13) in K562 cells. **A)** Schematic representation of target regions for CRISPR/Cas9-directed cleavage on chromosomes 7 and 12 used for guide RNA (gRNA) design. Fusion junctions are flanked by arrows representing PCR primers used for confirmation, yielding products of 614 bp and 937 bp. Below, detailed location of the targeted regions on chromosome 7q36.3 and 12p13.2 with reference to known t(7;12) breakpoints used to generate the translocation. **B)** FISH using the specific t(7;12) probe XL t(7;12) MNX1/ETV6 (MetaSystems Gmbh, Altlussheim, Germany, **Supplementary Table 3**) hybridising chromosome 7q36 in red and chromosome 12p13 in green. Two yellow fusion signals, pointed by arrows, indicate the presence of t(7;12) in a representative interphase nucleus of K562-t(7;12). The karyotype of K562 is nearly tetraploid and harbours complex rearrangements, including a duplication of the 7q36 locus within the short arm of chromosome 7, hence showing five 7q36 signals and two 12p13 signals (30) (**Supplementary Figure 2**). **C)** Metaphase spread of K562-t(7;12) hybridised with the same probe shows the presence of derivative chromosomes der(7) and der(12), pointed by arrows. **D)** Confirmation of the presence of t(7;12) in K562-t(7;12) but not K562 control (‘ctrl’) cells (K562 electroporated with Cas9 only) by PCR amplification of fusion junctions and product separation on agarose gel; bands correspond to the predicted sizes shown in panel A. NTC = no template control. **E)** qRT-PCR validation of overexpression of *MNX1* in K562-t(7;12) compared to K562 control. The fold change was calculated using the ΔΔCt method by normalisation to the endogenous gene *HPRT1*. Error bars represent standard deviation (SD) of n=3. Primers are reported in **Supplementary Table 4. F)** A custom-made 3-colour probe (MetaSystems dual-colour ETV6 + PAC-derived RP5-1121A15) hybridising *ETV6* portions in red (centromeric) and green (telomeric), and the *MNX1* locus in cyan (**Supplementary Figure 2; Supplementary Table 3**), allowed visualisation of both derivative chromosomes in interphase nuclei (pointed by yellow arrows). The white radius arrow represents the distance between nuclear interior (value=0) and nuclear periphery (value=1), which was used in the calculation of radial nuclear locations (RNL). **G)** RNL of der(7) and der(12) signals in K562-t(7;12) interphase nuclei. The RNL values are expressed as median values of individual localisations of FISH signals (200 nuclei analysed per condition). Errors bars represent standard error of the mean (SEM). Values closer to 0 indicate an internal position within the nucleus (described in detail in Federico *et al*. (31)).

While the transformed nature of the K562 cell line did not allow assessment of the transforming capacity of t(7;12) in this system, we observed a distinct phenotype associated with the K562-t(7;12). In clonogenic colony-forming assays on methylcellulose-based medium, K562-t(7;12) produced a significantly higher proportion of colonies with a diffuse phenotype, at the expense of the more prevalent compact colonies of K562 control (**Figure 2A-B**). K562 produces mostly erythroid-like compact colonies, while the diffuse phenotype associates with an immature myeloid identity (CFU-G or CFU-GM-like) (21), suggesting that t(7;12) changes the erythroid differentiation bias of K562. Importantly, the phenotype was maintained through replating, indicating that the translocation can be perpetuated through self-renewal. This is in contrast with the effects of *MNX1/ETV6* and *MNX1* overexpression in adult haematopoietic tissues, for which phenotypic changes could not be maintained through self-renewal (10,13).

**Figure 2.**
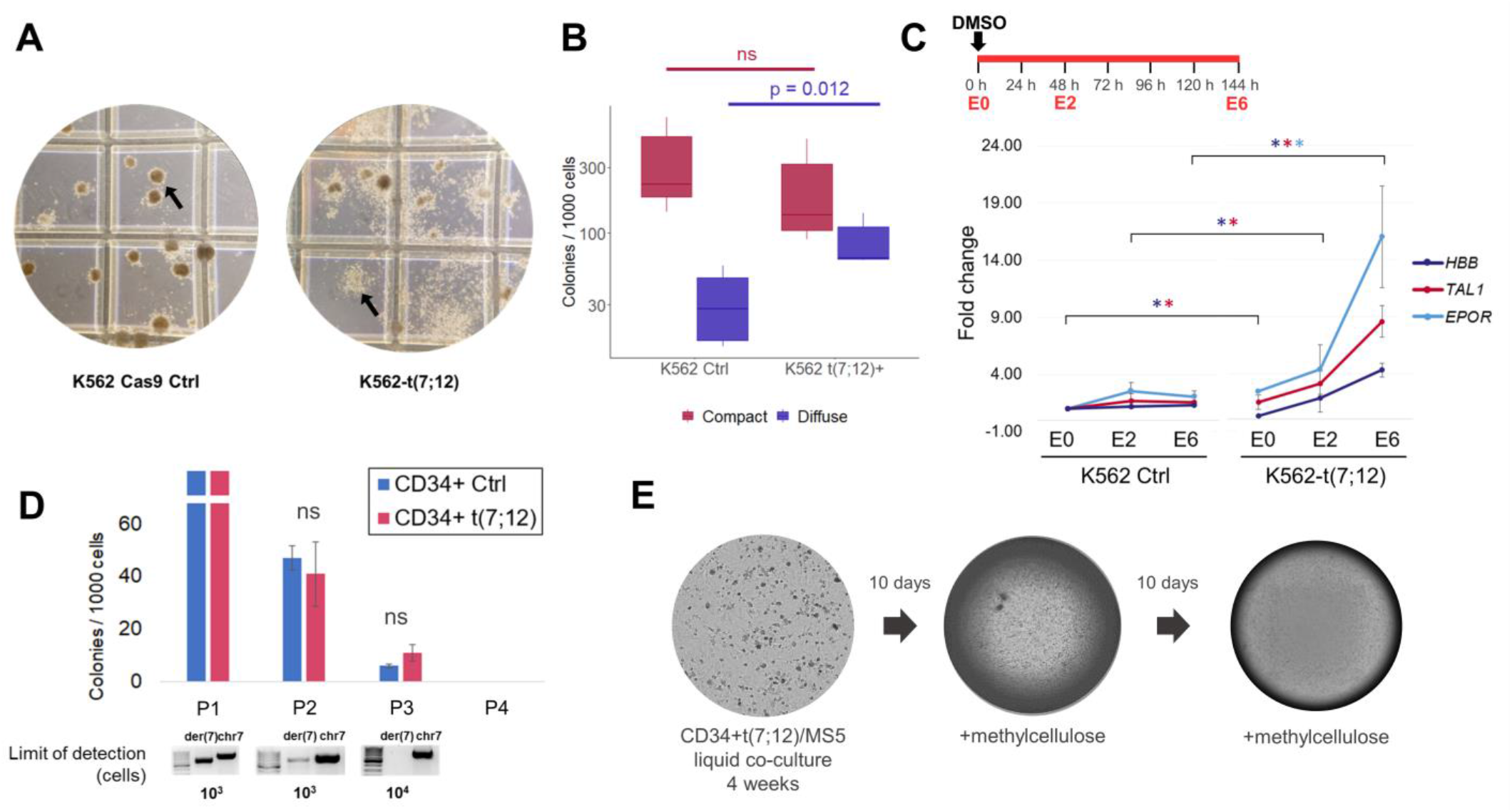
Functional effects of t(7;12) in haematopoietic cells. **A)** Representative images of colony-forming assays of K562 control (‘K562 Cas9 Ctrl’) (left) and K562-t(7;12) (right) on Methocult H4434 Classic (StemCell Technologies). Arrows indicate distinct morphologies of compact (left) and diffuse (right) colonies. **B)** Frequency of colony formation per 1000 K562 Cas9 Ctrl and K562-t(7;12) plated cells. Mean and 95% CI of n=3 independent experiments. Statistical significance determined by 2-tailed Student’s t-test. **C)** Quantitative RT-PCR gene expression analysis of erythroid differentiation of K562-t(7;12) cells; schematic representation of differentiation assay (top left). 100 000 cells were plated in RPMI medium supplemented with foetal bovine serum (FBS) with 1.5 μM DMSO added at time point 0 h. ‘E’ indicates the day of erythroid differentiation, from E0 (0 h) to E6 (144 h). Expression of erythroid markers *EPOR, TAL1*, and *HBB* was quantified in K562-t(7;12) through E0 to E6, calibrated to K562-Ctrl E0. Fold changes were calculated using the ΔΔCt method by normalisation to the endogenous gene *HPRT1*. Asterisks indicate statistically significant fold changes of K562-t(7;12) compared to K562 control for each gene. Full statistical tests on all conditions are shown in **Supplementary Figure 3**. Primers are reported in **Supplementary Table 4. D)** Introduction of the t(7;12) translocation in healthy CD34+ haematopoietic progenitors enriched from mobilised peripheral blood by magnetic separation. Serial replating colony-forming assays in Methocult H4434 Classic medium with multi-lineage cytokines. 10 000 CD34+ HSPCs were plated in methylcellulose-based medium and allowed to form colonies. Colonies were scored, collected and dissociated after each plating, and replated over several rounds. HSPCs are expected to originate progressively less colonies, unless a survival advantage is gained through transformation. ‘P0-P4’ indicate the order of plating. Progressively less colonies were produced, without statistically significant differences between the two conditions. Agarose gel electrophoresis of semi-quantitative PCR is shown underneath each colony count, indicating the limit of detection of the der(7) fusion junction. A non-translocated region of chromosome 7 was used as a control (‘chr 7’) to confirm the presence of amplifiable template DNA. **E)** MS5 stroma with the addition of CD34+t(7;12) cells. Two colonies were visible following methylcellulose addition after 4 weeks of co-culture. No colonies visible following a second round of co-culture and methylcellulose addition.

We further confirmed the depletion of erythroid identity by subjecting both K562-t(7;12) and K562 control to induced erythroid differentiation assay by adding DMSO to the culture medium and assessing the expression of erythroid marker genes (22) (**Figure 2C; Supplementary Figure 3**). At the beginning of the assay at erythroid day 0 (E0), we observed a lower expression of *HBB* and *TAL1* in K562-t(7;12) compared to K562 control (**Supplementary Figure 3A**), indicating an attenuation of erythroid signatures, compatible with the myeloid bias of K562-t(7;12) colony formation. Control K562 underwent differentiation and plateaued at E2, as seen by upregulation of *HBB, TAL1*, and *EPOR*. In K562-t(7;12), the differentiation capacity was not blocked by the presence of the translocation as seen by the upregulation of the three genes, albeit displaying higher fold changes compared to the control (**Figure 2C; Supplementary Figure 3**). In this light, the erythroid differentiation assay confirmed that the erythroid differentiation programme was still achievable in K562-t(7;12), but at higher fold gene expression changes, suggesting retainment of erythroid cells at an earlier differentiation state in the absence of external differentiation cues, and/or relative expansion of myeloid-affiliated cells as a result of the translocation. Together with the colony forming capacity suggesting an imbalance between erythroid and granulocytic/monocyte-like colonies (**Figure 2A**), we demonstrate that t(7;12) can affect erythroid differentiation programmes. In earlier models of t(7;12), *MNX1*-overexpressing bone marrow cells transplanted into recipient mice were primarily found in myeloid-committed progenitor populations and particularly within megakaryocytic/erythroid progenitors (13). In human cord-blood CD34+ HSCPs, *MNX1* overexpression induced erythroid transcriptional programmes and reduced overall colony-forming capacity (13), indicating that t(7;12) encompasses but may exceed the observed effects of *MNX1* overexpression on erythroid differentiation.

In order to understand t(7;12) effects in untransformed haematopoietic cells, we introduced the translocation in adult CD34+ HSCPs. Despite being able to confirm its presence by PCR (**Supplementary Figure 4A**), the frequency of t(7;12)-harbouring cells decreased through serial replating in methylcellulose-based colony-forming assays (**Figure 2D**), as well as in liquid culture (**Supplementary Figure 4B**), as determined by semi-quantitative PCR. We used Long-Term Culture Initiating Cell (LTC-IC) assays to test for leukaemia-initiating effect. This 2-step assay involves a 4-week co-culture with MS5 stroma, which selects for self-renewing or slowly proliferating HSPC, followed by assessment of HSPC functional potential in colony-forming assays. Similarly to short-term clonogenic assays, edited CD34+ cells did not show any advantage in persistence of LTC-IC, as measured by the lack of sustained colony formation following co-culture in MS5 and serial methylcellulose replating (**Figure 2E**). Taken together, our observations are in line with the previously reported inability of *MNX1* overexpression to transform adult haematopoietic cells (10,13). This may suggest a requirement for additional genetic events to achieve full *MNX1*/t(7;12)-associated transformation, or otherwise suggest that t(7;12) selectively affects a cell type or developmental stage not efficiently represented in adult mouse or human haematopoietic tissues. The transformed nature of K562 cells, with or without contribution from the persistence of foetal developmental programmes, as evidenced by the nature of haemoglobins expressed (23), may provide a more suitable substrate for persistence of t(7;12).

In order to understand if K562-t(7;12) recapitulated molecular signatures identified in t(7;12) AML patients, we performed RNA sequencing analysis of K562-t(7;12) compared to K562 control, and identified 436 upregulated and 116 downregulated genes by p value lower than 0.01, of which 196 under the threshold of 1% false discovery rate (FDR) (**Figure 3A**). Gene Ontology (GO) analysis revealed that differentially expressed genes (filtered by FDR) were involved in functions related to cell adhesion, immune response, and transport-related biological processes (**Figure 3B**). Genes associated with cell adhesion and the structural component of the matrix included collagen genes (*COL*), integrins (*ITG*), and the surface markers CD24 and CD84, which were also identified by further dissection of the upregulated genes by molecular function (**Figure 3D**). The upregulation of genes associated with cell adhesion and cell-cell interactions was previously reported by Wildenhain *et al*. (10), from the differential expression of adhesion-related genes in t(7;12) patients compared to *MLL* patients (i.e. *EDIL3, CNTNAP5, ANGPT1, DSG2, ITGA9, ITGAV, KDR*, and *SIGLEC6*). This gene expression profile was highly consistent with K562-t(7;12) (50% of upregulated genes matching), including an upregulation of contactin genes *CNTN1* and *CNTN4, ANGPT1*, a number of integrins (*ITGA2B, ITGA4, ITGA8, ITGA9, ITGAM, ITGAX, ITGB2*), and sialic acid-binding immunoglobulin-type lectins (*Siglecs; SIGLEC6, SIGLEC10, SIGLEC14, SIGLEC17P, SIGLEC8, SIGLEC22P*) (**Figure 3C; Supplementary Figure 5**). Indeed, the involvement of cell adhesion and extracellular matrix modelling in bone marrow niche interactions has been proposed as an important leukaemic mechanism of t(7;12) (10,12,13,24). Our data capture a number of these genes and can provide a model for future studies of individual and coordinated gene contributions to t(7;12) biology. The signature from Wildenhain *et al*. (10) also included the downregulation of *HOXA* genes in t(7;12) compared to *MLL* patients, which we did not find as differentially expressed in K562-t(7;12) (**Supplementary Figure 5**). This reflects the specificity of *HOXA* dysregulation in *MLL* leukaemia (25), but not necessarily the relevance in t(7;12). Balgobind *et al*. (26) also described a distinctive gene expression signature of t(7;12) leukaemia compared to a broader set of AML subtypes including *MLL* rearrangements, t(8;21), inv(16) and t(15;17), identifying 12 discriminative genes (*TP53BP2, DNAAF4, EDIL3, LIN28B, BAMBI, MAF, FAM171B, AGR2, CRISP3, CTTNBP2, KRT72*, and *MMP*). Our model matched a differential expression of *TP53BP2, LIN28B, CTNNBP2*, and *MPP2*, with half of this signature with matched or partially matched genes (**Figure 3C; Supplementary Figure 5**). Downregulated genes were enriched for molecular functions of macromolecule binding (**Figure 3E**). Several haemoglobin genes were found to be downregulated (*HBZ, HBG1-2, HBA1-2, HBB*, and *HBE1*) (**Supplementary Figure 6**), which are related to as functional categories of haptoglobin, haemoglobin, and oxygen binding processes (**Figure 3E**). The concomitant downregulation of peroxiredoxins (*PRDX1, PRDX4*, and *PRDX6*) also highlighted peroxidase and antioxidant activities, which are consistent with biological functions of erythrocytes, and therefore accompany the relative depletion of erythroid identity (**Figure 2A-C**).

**Figure 3.**
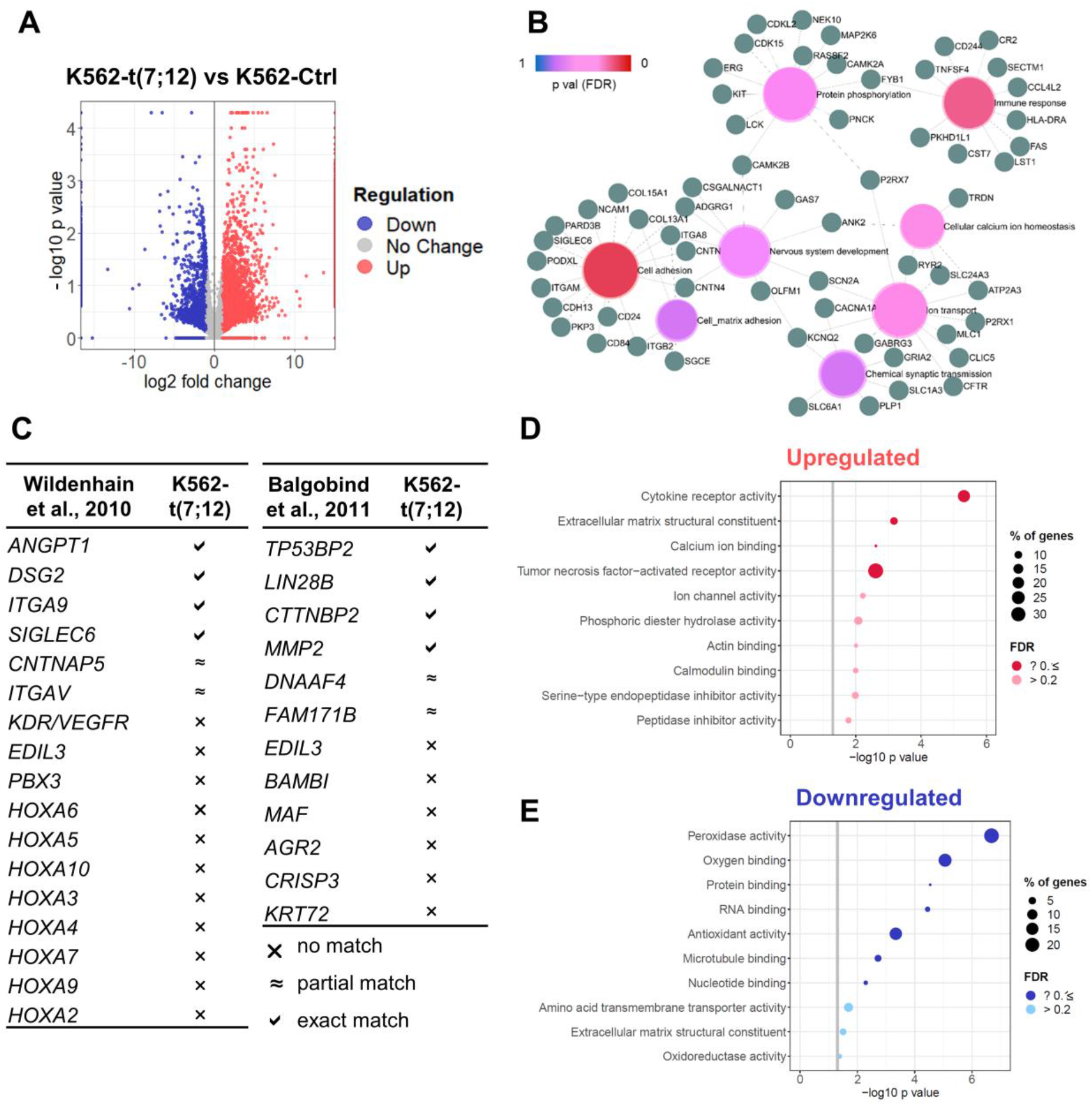
Transcriptional analysis of K562-t(7;12). **A)** Volcano plot showing differentially expressed genes in K562-t(7;12) compared to K562 control, depicting up- (in red) and downregulated (in blue) genes and their statistical significance by -log10 p value. Genes with an absolute log2 fold change lower than 1 are shown in grey (‘No change’). **B)** Gene Ontology network plot of differentially expressed genes by false discovery rate (FDR) < 0.1 showing significant biological processes (PANTHER) and genes associated to each GO term, constructed on ExpressAnalyst (available at www.expressanalyst.ca) **C)** Comparison of differentially expressed genes in K562-t(7;12) and published reports of genes associated with t(7;12) patients. Symbols indicate difference in expression between K562-t(7;12) and K562 control by statistical significance (threshold p ≤ 0.05), defined as an exact gene match, a partial match (i.e. dysregulation of a gene belonging to the same family), or no match. **D-E)** GO terms relating to molecular functions (PANTHER) of upregulated **(D)** and downregulated **(E)** genes in K562-t(7;12). The grey intercept marks the p=0.05 threshold in –log10. The size of the dots indicates the percentage of genes mapping to a specific term. Dark red or blue indicates an enrichment with an FDR ≤ 0.1.

Overall, observed gene expression changes were consistent with individual gene signatures reported in analyses of t(7;12) patients (**Figure 3C**). We sought to further validate the clinical relevance of the K562-t(7;12) model by a systematic comparison with microarray and RNA sequencing data of t(7;12) patients from Balgobind *et al*. (26) and the TARGET database (28). We derived a unique 177-gene signature by comparison of t(7;12) with other individual paediatric subtypes (inv16, MLL rearrangements, normal karyotype = NK, t(8;21), and other) and identification of the common differential gene set (**Figure 4A**). Similarly, we defined a 122-gene list uniquely differentially expressed between t(7;12), but not other AML subtypes, and normal paediatric bone marrow (**Figure 4B**). Consistent with previous reports (10), both gene sets were enriched for cell adhesion (collagen genes, *SIGLEC8*, contactins), cellular transport and lipid metabolism gene ontologies (**Figure 4C-D**). We also note that *HOXA* genes, previously identified by comparing t(7;12) AML with *MLL* rearrangements (10), were not present in these signatures as a result of comparisons with multiple paediatric AML subtypes. Importantly, we used the 177 and 122 t(7;12) genes as custom gene sets for Gene Set Enrichment Analysis (GSEA), and showed significant enrichment in the K562-t(7;12) model (**Figure 4E-F**), thus demonstrating that engineering of the translocation in K562 cells resulted in a clinically-relevant model. Extraction of the core enriched genes, or leading edge of GSEA identifies individual patient-specific genes recapitulated in our model (**Figure 4G**), which can be regarded as future targets in mechanistic studies of t(7;12)-associated phenotypes. Crucially, *MNX1* was identified as a leading-edge gene, asserting its centrality to t(7;12) K562 biology (**Figure 4G**).

**Figure 4.**
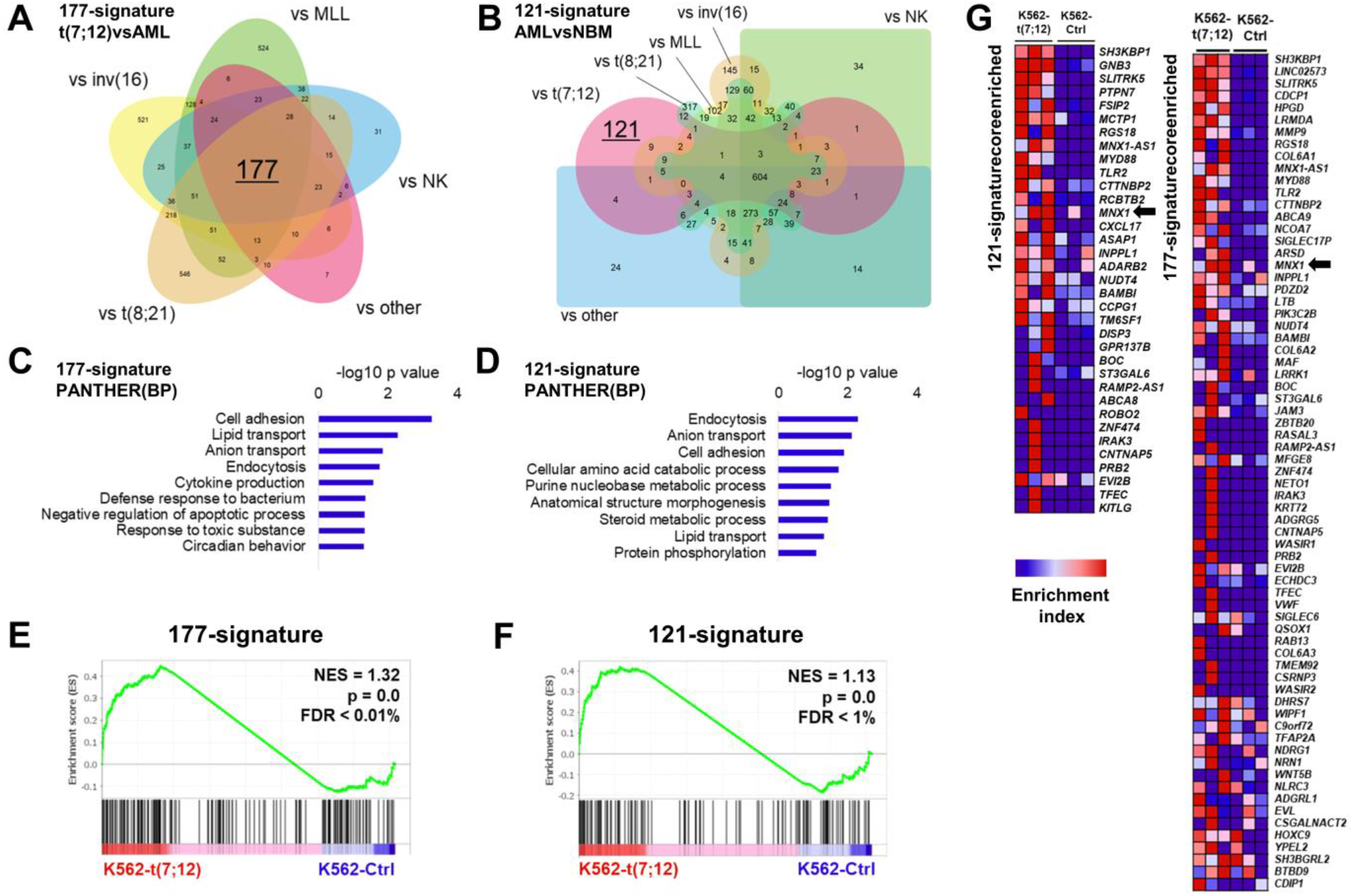
Comparison of K562-t(7;12) transcriptional landscape with t(7;12) patient signatures. **A)** 177-gene signature of t(7;12)-patient gene expression extrapolated from published microarray and RNA sequencing datasets by comparison with other paediatric AML subtypes. The Venn diagram shows the 177 intersect genes. **B)** 121-gene t(7;12)-specific signature inferred by comparisons of gene expressions of paediatric AML and normal bone marrow (NBM) samples. Edward’s Venn diagram highlights the 121 exclusive genes to t(7;12). **C-D)** Gene Ontology analysis of the 177-signature and 121-signature by the PANTHER annotation repository of biological processes (BP). **E-F)** GSEA enrichment plots of K562-t(7;12) gene expression profile using the 177- and 121-signatures. NES = normalized enrichment score; FDR = false discovery rate. **G)** Core enriched genes from the GSEA using the 177- and 121-signatures, shown by their enrichment index in K562-t(7;12) against K562 control. *MNX1* is highlighted by the arrows.

During the preparation of this manuscript, another group successfully recreated the t(7;12) translocation in human induced pluripotent stem cell (iPSC) using CRISPR/Cas9 (29), with comparable features to our model. The authors observed an increased frequency of erythroid and myeloid progenitors in colony forming assays, coherent with an increase in myeloid transcriptional programmes. Notably, the iPSC model reproduced the overexpression of *MNX1* and production of a *MNX1/ETV6* fusion transcript, albeit at a low transcriptional level that questions its biological relevance. Despite extensive FISH analyses in early studies (4-6), the exact breakpoints required to produce the fusion transcripts from t(7;12) have not been elucidated. In Nilsson et al. (29), the expression of the *MNX1/ETV6* mRNA was achieved by targeting a similar genomic region to us on 7q36, but a distinct intronic region of *ETV6*. In our K562 model, a chimaeric transcript was not detected by potential gene fusion search by TopHat-Fusion on the RNA sequencing reads; an interesting observation given the 50% occurrence of chimaeric transcripts in patients (2). With the continued development of genome editing technologies, we foresee that these *in vitro* models will allow a more precise dissection of the breakpoint regions required for the formation of the yet uncharacterised *MNX1/ETV6* chimaera. Both the iPSC and K562 models of t(7;12) showed similarities to patient-specific gene expression patterns, which strengthens the validity of CRISPR/Cas9 to generate disease models based on clinical genomic information.

In summary, we generated cellular model of t(7;12) translocation which recapitulates *MNX1* overexpression and captures cell adhesion signatures putatively core to t(7;12) oncogenic programmes. The model displays a granulocytic-monocytic phenotype compatible with the lineage affiliation of clinical t(7;12) AML, and disrupts erythroid lineage programmes as described in *MNX1* overexpression. We confirmed the inability of t(7;12) to transform adult haematopoietic tissues, highlighting the requirement for additional oncogenic hits, or a specific cellular background which may explain the unique association of t(7;12) with infants. Complementarily to earlier *MNX1*-based models (10,13), and more recent genome edited tools (29), our t(7;12) K562 model will provide a framework for the molecular elucidation of this unique subtype of infant AML.

## Acknowledgements

Dr Giorgia Santilli is acknowledged for providing adult CD34+ cells. DR is the recipient of a Kidscan funded PhD studentship and partly supported by Brunel University London. YC and AS are supported by a grant from the Oracle Cancer Trust.

## Author Contributions

DR was responsible for carrying out all experiments, extracting and analysing data, interpreting results, producing tables and figures and writing the manuscript. YC performed FISH experiments, captured and analysed fluorescent microscopy images, participated in editing and reviewing the manuscript. CF, SS, and FB contributed the gene positioning analysis and provided feedback on the manuscript. RS participated in gene expression analysis of K562 erythroid differentiation. CS supervised the computational analysis of transcriptomic data and reviewed the manuscript. CP designed and supervised functional analysis of K562 and CD34+ cells, participated in design and interpretation of transcriptomic data analysis, and reviewed the manuscript. AS was responsible for designing and supervising the project, with specific input on the gene editing, and reviewed the manuscript. ST was responsible for designing and supervising the project, with specific input on the chromosome biology aspects including interpretation of FISH results, and reviewed the manuscript. All authors have read and agreed to the published version of the manuscript.

## Data Availability Statement

The RNA sequencing datasets generated and analysed during the current study have been deposited in the ArrayExpress repository under accession number E-MTAB-11851. The results published here are partly based upon data generated by the Therapeutically Applicable Research to Generate Effective Treatments (TARGET) (https://ocg.cancer.gov/programs/target) initiative, of the Acute Myeloid Leukemia (AML) cohort phs000465. The data used for this analysis are available at https://portal.gdc.cancer.gov/projects.

## Supplementary Figures

**Supplementary Figure 1.**
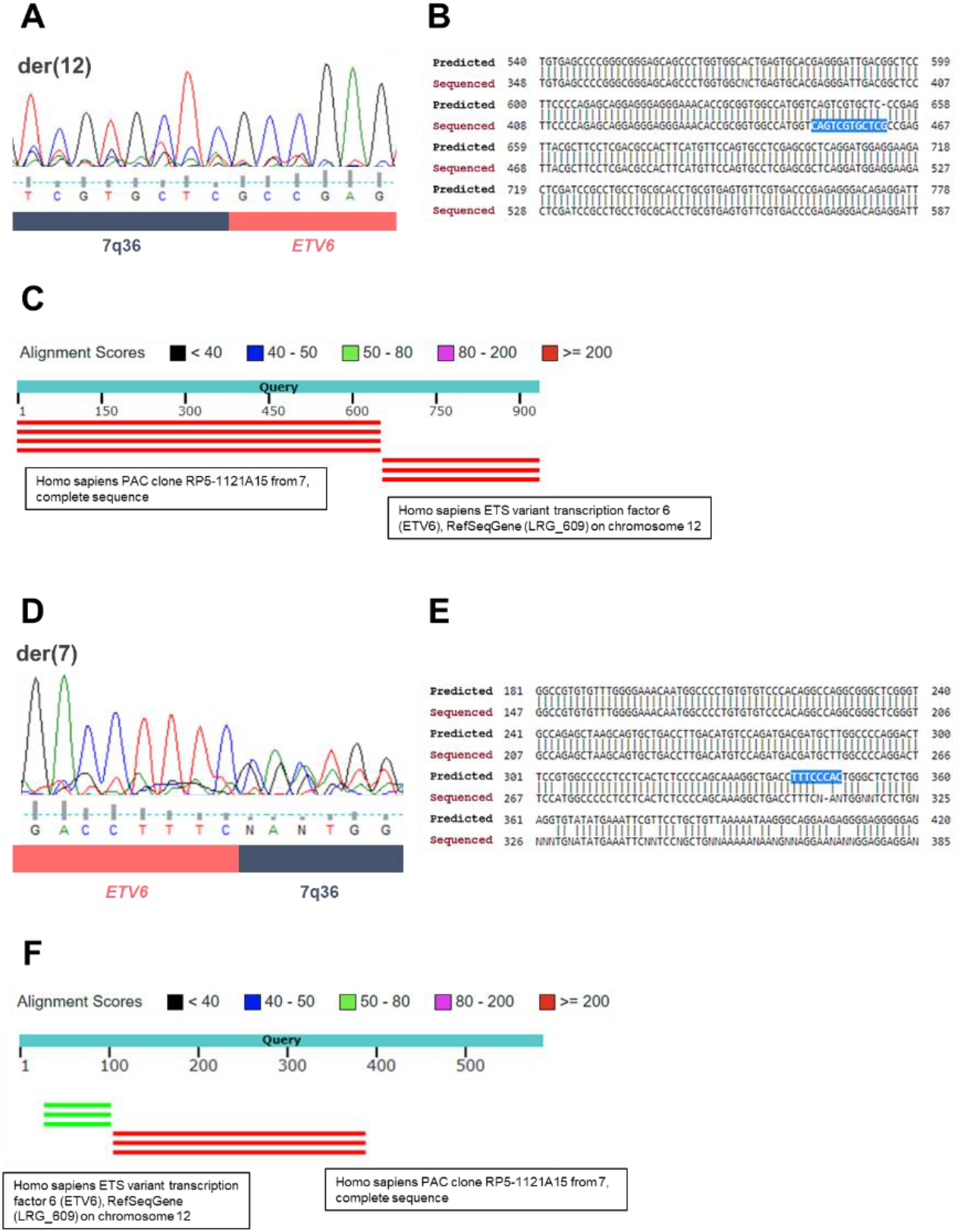
Sanger sequencing of der(7) and der(12) fusion junctions. Chromatograms showing the nucleotide sequence of amplified fusion junctions in der(12) **(A)** and der(7) **(D)**. Sequence alignments of der(12) **(B)** and der(7) **(E)** corresponded to the predicted sequence. Highlighted nucleotides in blue correspond to the fusion junction. The identity of the sequences was confirmed by BLAST, showing matching sequence identity from chromosome 7q36 region (RP5-112A15) and the ETV6 gene in both derivatives **(C**,**F)**.

**Supplementary Figure 2.**
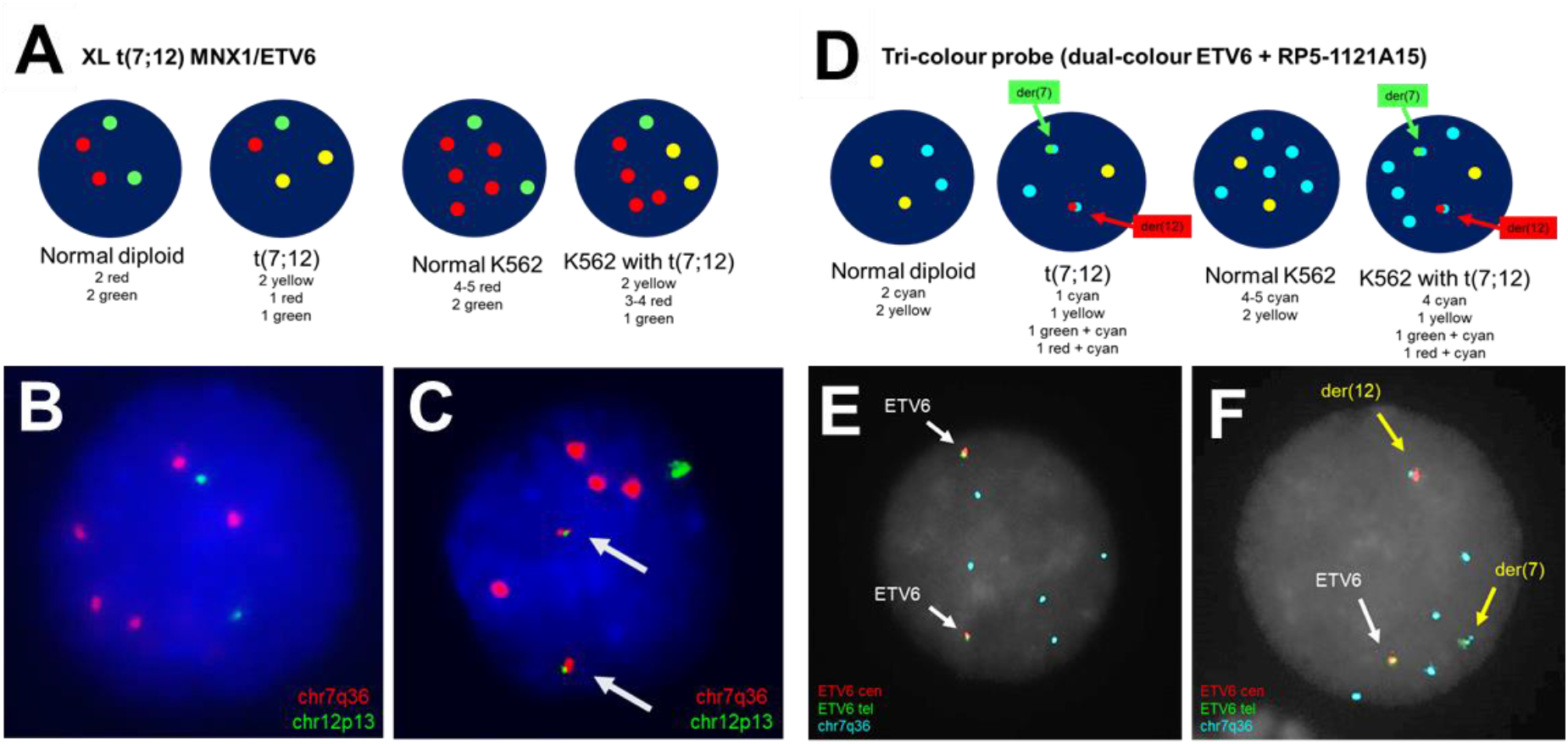
FISH on K562-t(7;12) and K562 control nuclei using two probe sets. **A)** The t(7;12)-specific FISH probe XL t(7;12) MNX1/ETV6 (MetaSystems) patterns in normal diploid nuclei and t(7;12) on the left, and normal K562 and K562 with t(7;12) on the right. The presence of the t(7;12) is confirmed by two yellow fusion signals, by splitting of the red and green signals due to the translocation. The karyotype of K562 is nearly tetraploid and harbours complex rearrangements, including a duplications of the 7q36 locus containing MNX1 within the short arm of chromosome 7, hence showing five 7q36 signals and two 12p13 signals (1) **B)** A normal pattern indicates the expected signal pattern in non-edited K562, with 5 red signals corresponding to the 7q36 locus, and 2 green signals hybridising to 12p13. **C)** Two yellow fusion signals, pointed by arrows, are indicative of the presence of the t(7;12). **D)** Tri-colour probe (dual colour ETV6 probe + RP5-1121A15) patterns in normal diploid nuclei and t(7;12) on the left, and normal K562 and K562 with t(7;12) on the right. The two derivatives are distinguishable by the fusion of the green and cyan (der7), and red and cyan (der12). **E)** Representative nucleus and metaphase of K562 Cas9 Ctrl showing the hybridisation patterns of the 3-colour probe with non-translocated *ETV6* and 7q36 regions, indicated by 2 yellow fusion signals and 5 cyan signals, respectively. **F)** Representative K562-t(7;12) nucleus showing the presence of two derivative chromosomes indicated by yellow arrows, the non-translocated *ETV6* allele (white arrow), and 4 non-translocated 7q36 signals. The der(7) is characterised by the fusion of the green and cyan signals, while the der(12) is distinguishable by red and cyan overlapping signals.

**Supplementary Figure 3.**
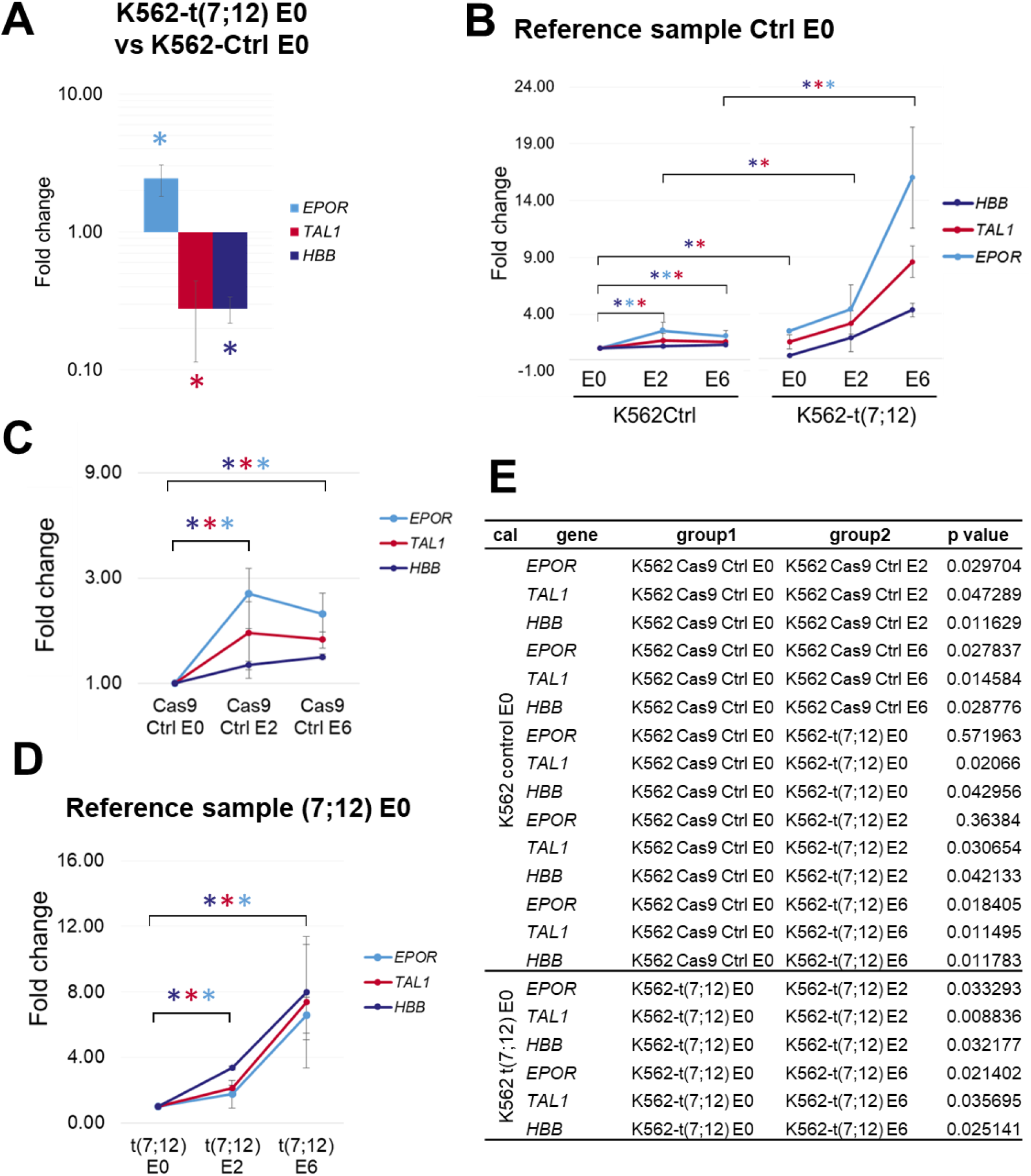
Erythroid differentiation assay of K562-t(7;12). **A)** Gene expression quantification of HBB, TAL1 and EPOR in K562-t(7;12) expressed in fold change compared to K562-Ctrl at the beginning of the erythroid differentiation assay (0 h; E0). Asterisks indicate a p value ≤ 0.05 with the colour corresponding to the gene as shown in the legend. Error bars indicate standard deviation on n=3. **B)** Full comparison of statistically significant gene expression changes, by fold changes calibrated to K562 control E0. **C)** Zoomed in view of K562 control with expanded fold change y axis. **D)** Fold changes in K562-t(7;12) calibrated to K562-t(7;12) E0, instead of K562 control. **D)** Table of p values determined by Student’s T-test in all combinations for expression changes across E0 – E2 – E6. “Cal” refers to the calibrator/reference sample of choice, either K562-Ctrl E0 or K562-t(7;12) E0. Group 1 and group 2 indicate the two conditions compared.

**Supplementary Figure 4.**
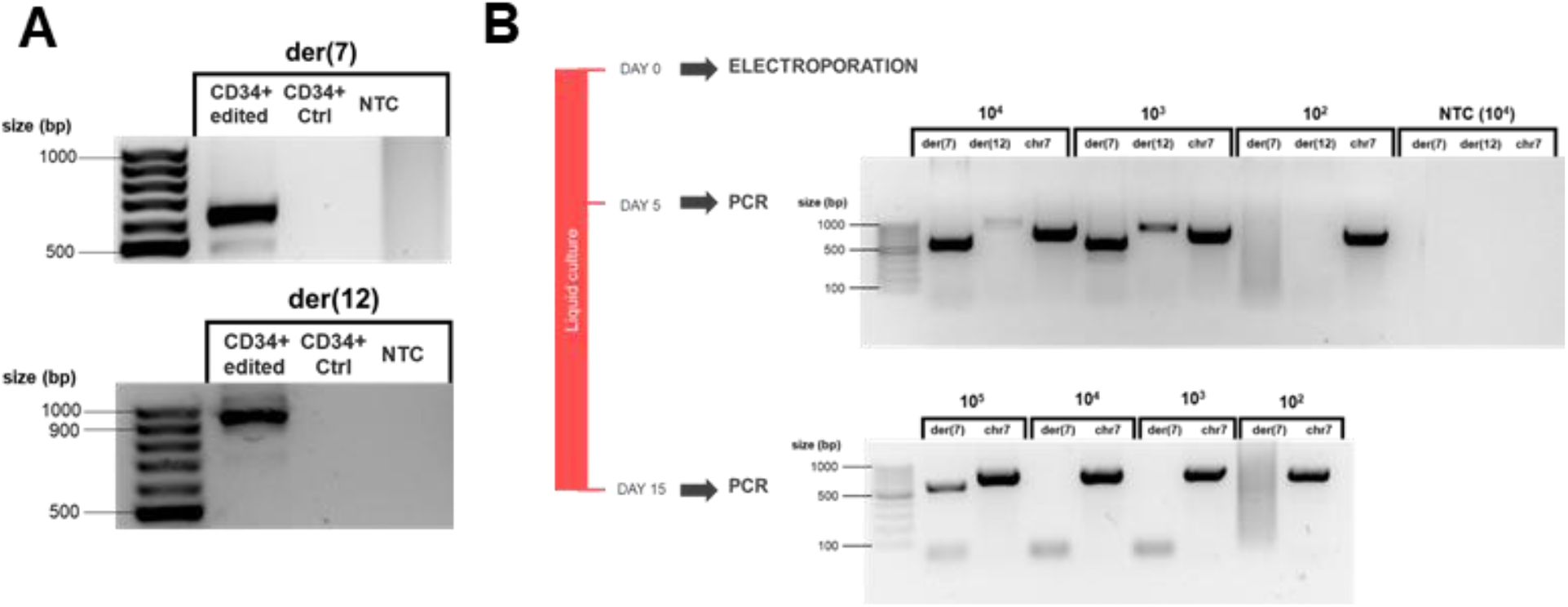
PCR amplification of fusion junctions of t(7;12) in edited CD34+ HSPCs. Agarose gel electrophoresis of amplified fusion junctions of der(7) (expected size 614 bp) and der(12) (expected size 937 bp) Ctrl = Cas9-only Control; NTC = no template control. Molecular marker: Gene Ruler 100 bp. Following electroporation, edited CD34+ HSCPs were grown in liquid culture for 15 days and screened for the presence of t(7;12) fusion junctions by semi-quantitative direct PCR. Varying amounts of cells were directly lysed and used as template for PCR to detect der(7) (614 bp) or der(12) (937 bp) fusion junctions. ‘Chr7’ indicates an unedited region of chromosome 7 used as control to confirm the presence of sufficient DNA template (735 bp). The lowest cell number at which a band is visible marks the limit of detection.

**Supplementary Figure 5.**
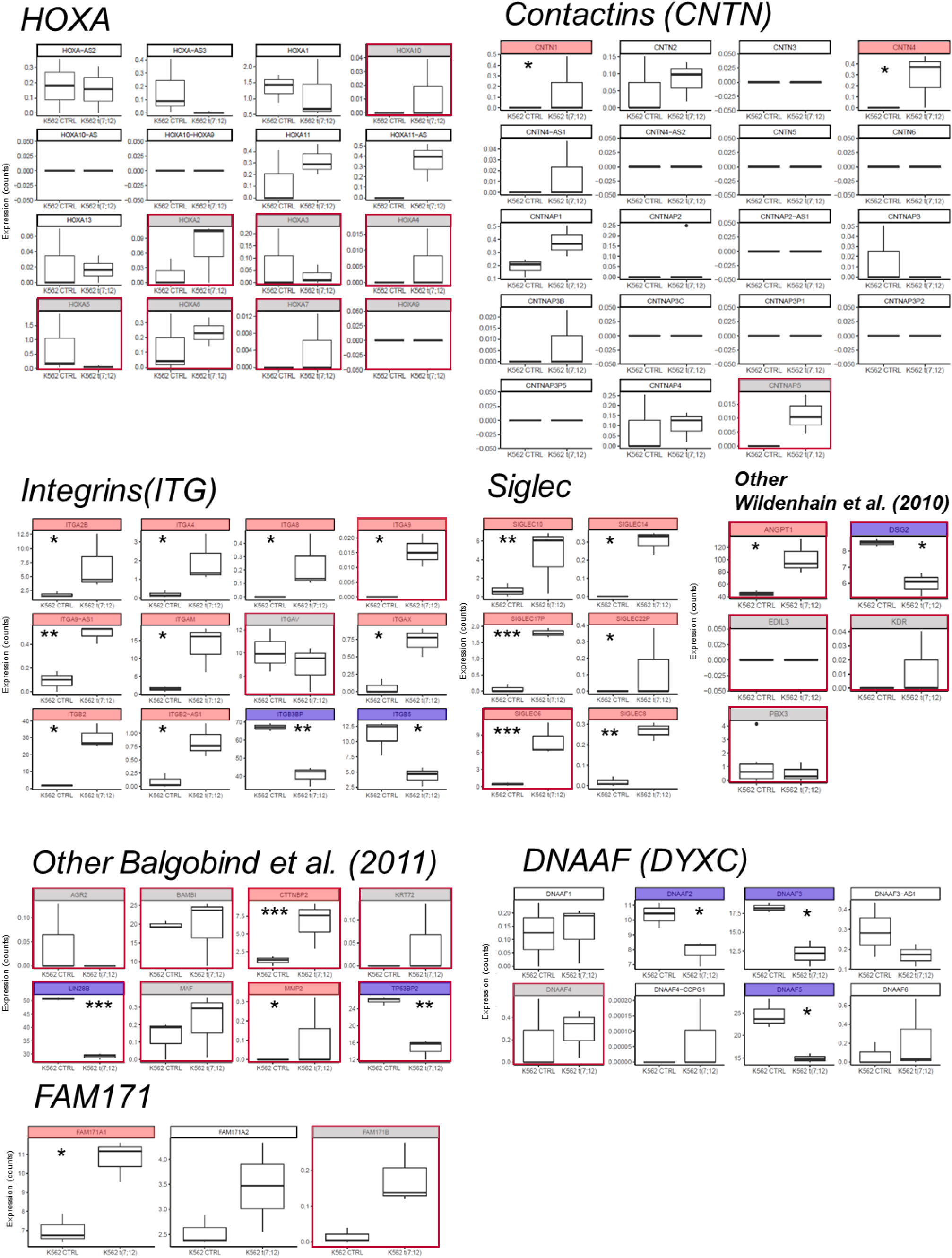
Expression differences in K562-t(7;12) compared to K562 control of known t(7;12)-associated genes. Gene expressions were extracted from RNA sequencing counts, and grouped by gene families. Red squares around the plot indicate a gene previously reported by Wildenhain *et al*. (2010) or Balgobind *et al*. (2011) as specific for the t(7;12) subtype. Grey shading indicates that the gene was not significantly dysregulated in K562-t(7;12) determined by T-test, while red indicates a significant upregulation, and blue a significant downregulation. Asterisks symbolise p value thresholds of 0.05 (*), 0.001 (**), 0.0001 (***), and 0.00001 (****).

**Supplementary Figure 6.**
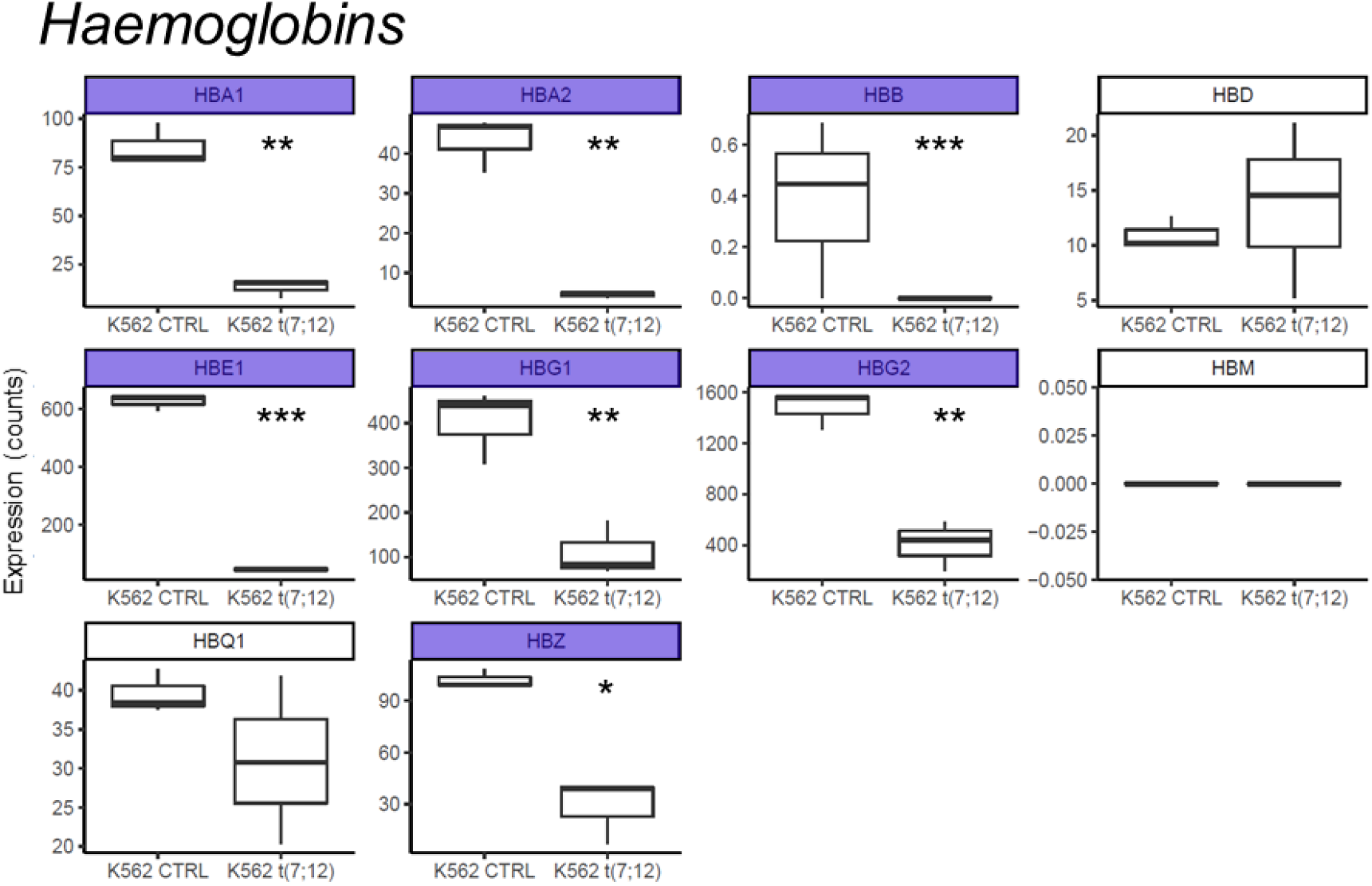
Differentially expressed haemoglobin genes in K562-t(7;12). Gene expressions were extracted from RNA sequencing counts of K562-t(7;12) and K562 control. Blue shading indicates a significant downregulation determined by T-test. Asterisks symbolise p value thresholds of 0.05 (*), 0.001 (**), 0.0001 (***), and 0.00001 (****).

## Supplementary Tables

**Supplementary Table 1.**
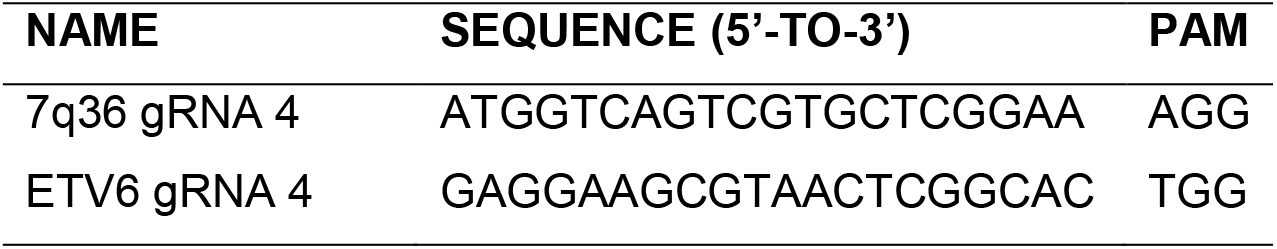
gRNA sequences.

**Supplementary Table 2.**
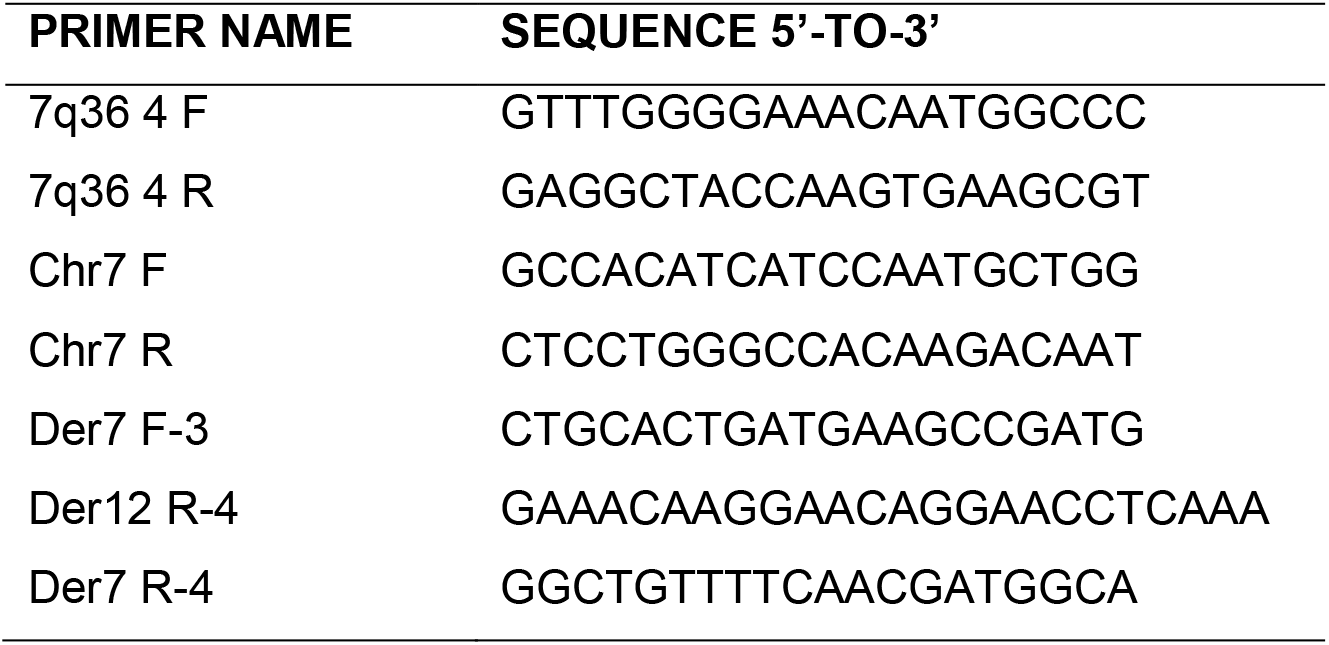
PCR primer sequences.

**Supplementary Table 3.** FISH probes ideograms.

| NAM<br>E | IDEOGRAM | FLUOROPHO<br>RES | MANUFACTU<br>RER |
| --- | --- | --- | --- |
| XL t(7;12) MNX1/ETV6 |  | FITC (ETV6)<br>Cy3 (MNX1) | Metasystems |
| Dual-colour<br>ETV6 |  | FITC (ETV6 5')<br>Cy3 (ETV6 3') | Metasystems |
| RP5-1121A15 |  | Cy5<br>(cyan false<br>colour) | PAC-clone |

**Supplementary Table 4.**
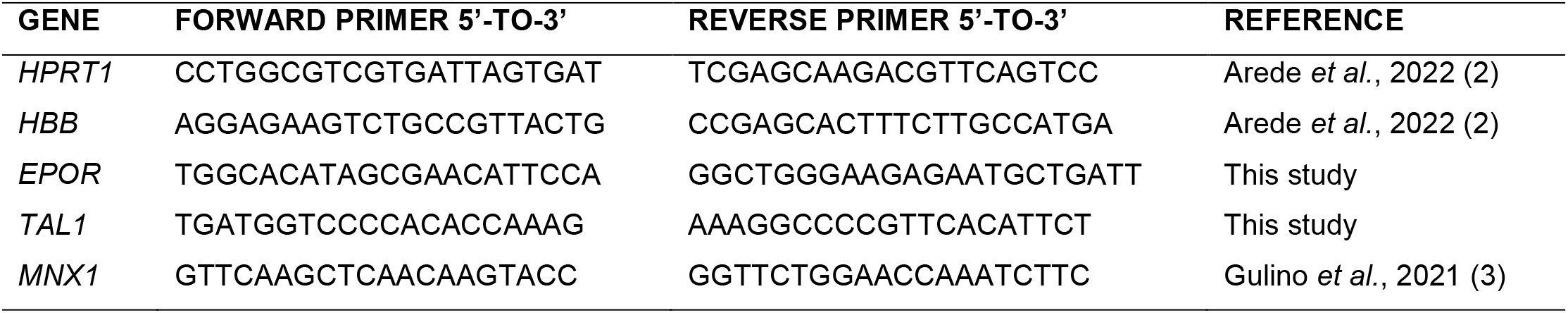
qPCR primer sequences.

**Supplementary Table 5.**
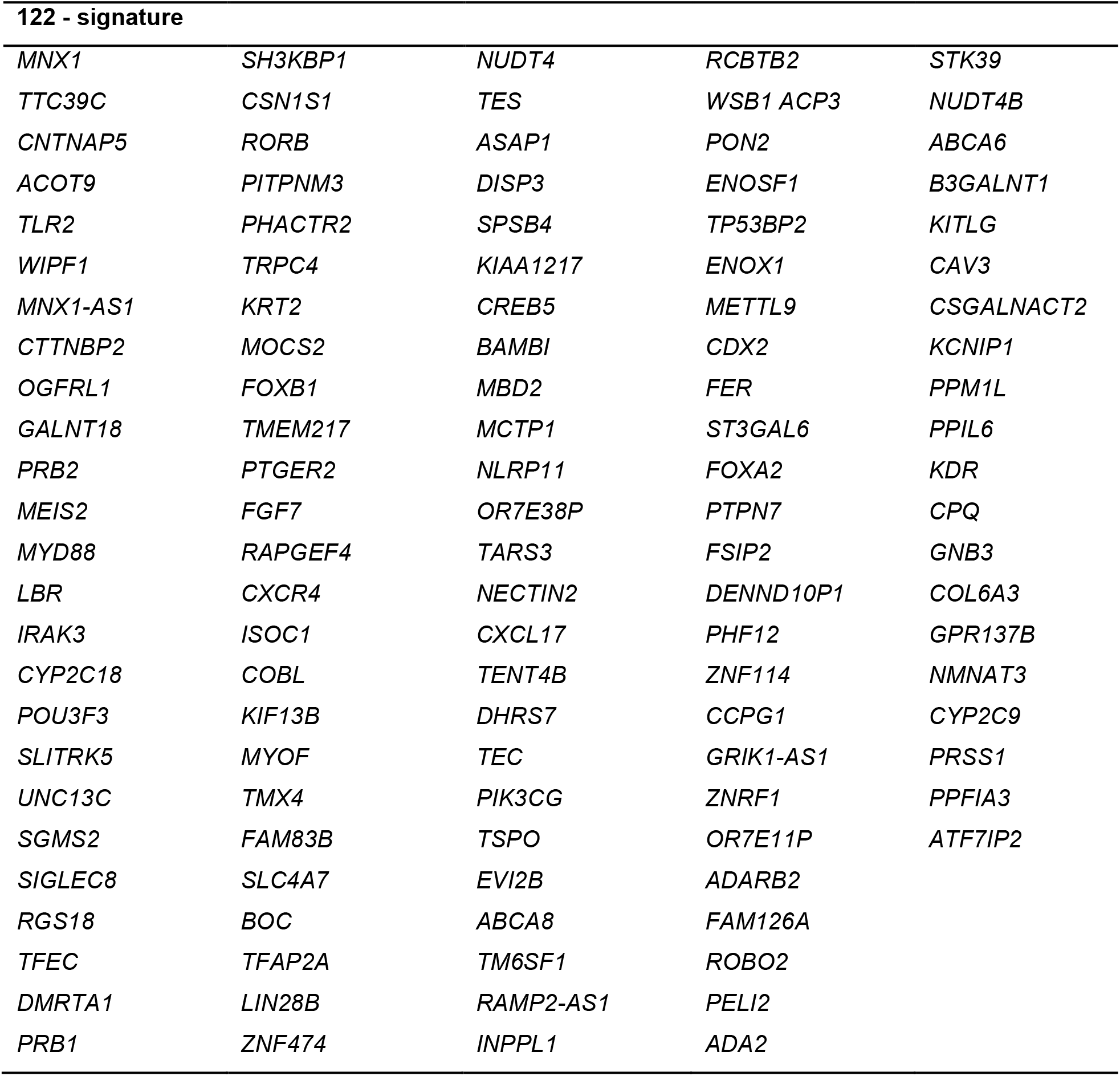
122-signature gene list.

**Supplementary Table 6.**
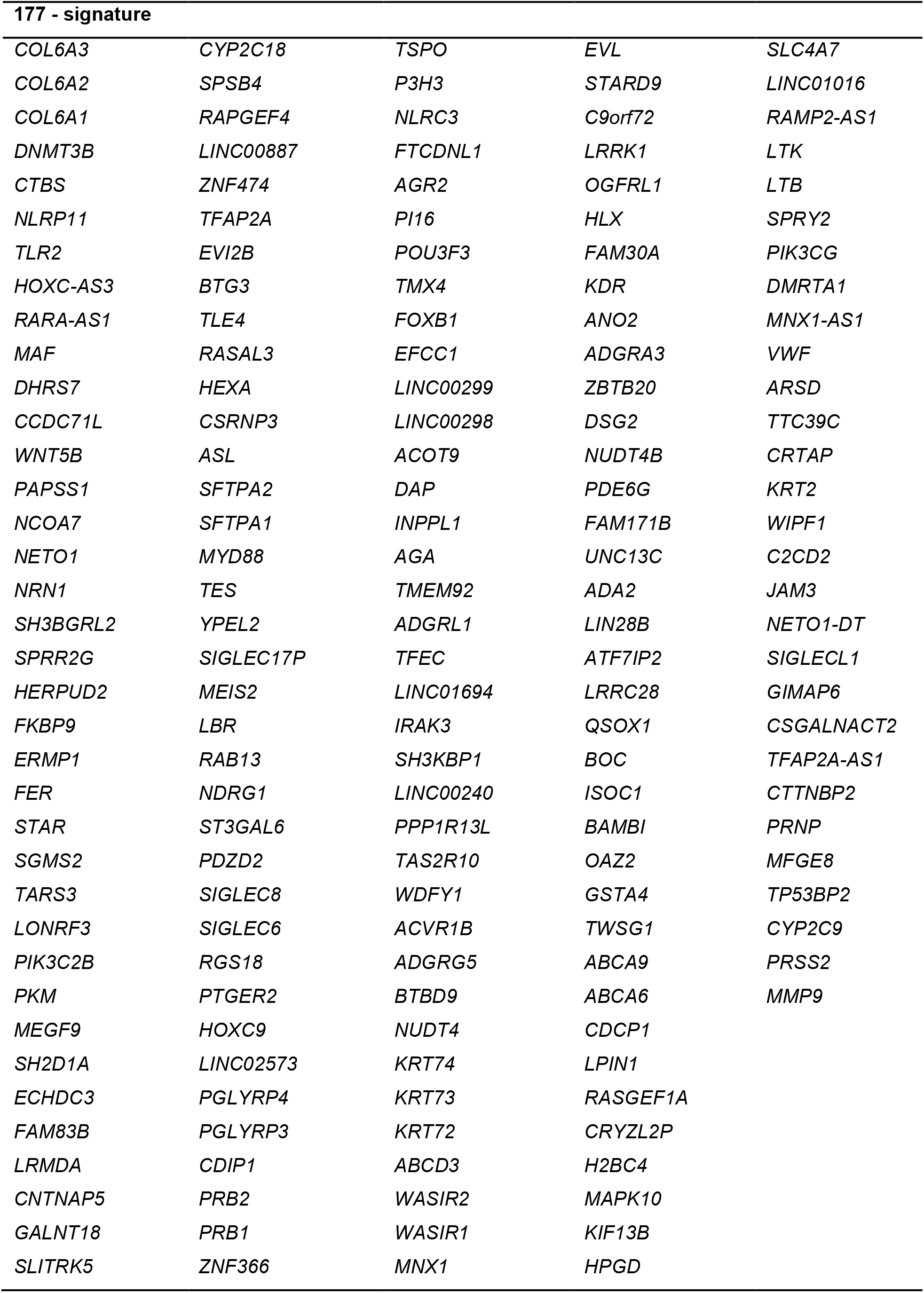
177-signature gene list.

## Methods

### 1. Cell cultures

The K562 leukaemia cell line was grown in RPMI1640 (Gibco, Paisley, UK), 10% foetal bovine serum (FBS) (Gibco) and 1% penicillin/streptomycin antibiotics (100 U/mL/100 μg/mL, Gibco). Peripheral blood mobilised CD34+ haematopoietic stem and progenitor cells (HSPCs) were obtained from UCL, Institute of Child Health, Infection, Immunity and Inflammation Programme (London, UK), purchased from commercially available apheresis products. CD34+ HSPCs were cultured in StemSpan medium (StemCell Technologies, Cambridge, UK) supplemented with 100 ng/ml SCF, 100 ng/ml FLT3-L, 20 ng/ml TPO, and 20 ng/ml IL-3 (Peprotech, London, UK). Cell lines were tested for Mycoplasma contamination using MycoSensor PCR Assay Kit (Agilent, Didcot, UK).

### 2. CRISPR/Cas9 editing

Sequences for human *MNX1* (accession number NC_000007.14) and *ETV6* (accession numbers NC_000012.12) were retrieved for gRNA design by the Invitrogen TrueDesign Genome Editor tool (threshold at 1 nucleotides of mismatch tolerance). gRNA sequences and the adjacent PAM sequence used in this study are reported in **Supplementary Table 1**. GeneArt Precision gRNA Synthesis Kit (Invitrogen, Inchinnan, UK) was used to synthesise customised gRNAs via in vitro transcription (IVT). The transfection of the gRNA and Cas9 endonuclease was achieved by assembly and delivery of an RNP complex via electroporation using the Neon Transfection System Kit (Invitrogen). The RNP complex was assembled at 1:2 molar ratio of TrueCut™ Cas9 Protein v2 (Invitrogen) to synthesised gRNA, and incubated at 37°C for 15 minutes prior to electroporation. Clones harbouring the translocation were isolated by single cell cloning by limiting dilution and PCR screening.

### 3. Amplification of t(7;12) fusion junctions by polymerase chain reaction (PCR)

PCR on genomic DNA was performed using High-Fidelity Phusion polymerase (Invitrogen) using manufacturer’s instructions. PCR primers (**Supplementary Table 2**) were designed using Primer-BLAST (NCBI). The complete genomic sequences of human MNX1 and ETV6 were obtained from the NCBI Gene and Nucleotide Database with accession numbers NC_000007.14 for *MNX1* and NC_000012.12 for *ETV6*. The amplified fragments of the correct estimated size for K562 Mut3 were excised, purified from the gels, and sequenced (**Supplementary Figure 1**).

### 4. Fluorescence in situ hybridisation (FISH)

Preparation of metaphase chromosomes was performed by colcemid treatment (0.05 μg/mL) as described (Federico *et al*., 2019) and fixed in methanol:acetic acid. FISH was performed using commercially available (MetaSystems, Altlussheim, Germany) and PAC-derived probes (**Supplementary Table 3**). PAC-derived probes were extracted from bacterial cultures and fluorescently labelled by nick translation (Roche, Mannheim, Germany). Following hybridisation and washes according to published protocols (Federico *et al*., 2019), images were captured using a Leica DM4000 fluorescence microscope (Leica Microsystems, Wetzlar, Germany). A minimum of 200 nuclei were captured for analysis. Representative images of K562 control and K562-t(7;12) nuclei hybridised with the probe combinations and the interpretation of signal patterns are shown in **Supplementary Figure 2**.

### 5. RNL analysis

Radial nuclear location (RNL) was calculated using 2D analysis of nuclei hybridised by FISH as previously described (4). The RNL is defined as a numerical value corresponding to the positioning of a FISH signal as the ratio of the nuclear radius. RNL ranges from 0 to 1, with 0 marking the interior and 1 marking the outer extreme. A minimum of 200 nuclei per condition are analysed; RNL is expressed taking into consideration the median value of all signals ± confidence interval, with a 0.650 and lower defining the nuclear interior (4).

### 6. Real-time quantitative PCR (qPCR)

Total RNA was extracted using the Monarch Total RNA Miniprep Kit (New England Biolabs, Hitchin, UK), from which complementary DNA (cDNA) was synthesised by reverse transcription using High-Capacity RNA-to-cDNA Kit (Applied Biosystems, Waltham, US). qPCR was performed using FastGene 2x IC Green Universal qPCR Mix (fluorescein) (Nippon Genetics, Düren, Germany) for each gene with primers listed in **Supplementary Table 4**. Differential gene expression was calculated using the ΔΔCt method. *HPRT-1* was used as endogenous reference gene.

### 7. Colony forming assay

Cells were plated onto semi-solid methylcellulose for colony-forming assays (CFC), in Methocult H4434 Classic (StemCell Technologies, Cambridge, UK). Cells were first suspended in IMDM (Gibco) + 20% FBS (Gibco) at the desired concentration (1000 – 10000 cells / plate). Plates were incubated at 37°C and 5% CO2 for 10-12 days, when colonies were scored.

### 8. Long-Term Culture Initiating Cell Assay (LTC-IC)

The MS5 stroma cells were plated in a 96-well plate at a density of 30 000 cells per plate. 45 000 CD34+ HSCPs were added to each well in 100 μl of H5100 Myelocult (StemCell Technologies), 1 μM hydrocortisone (StemCell Technologies), 20 ng/ml of IL-3, TPO, G-CSF, and SCF (Peprotech). The medium was replaced weekly and the assay was conducted for 4 weeks. At the end of the 4th week, 100 μl of Methocult H4434 (StemCell Technologies) was added to each well, and colonies were scored after 10 days. If colonies were present, the methylcellulose-based culture was dissociated by extensive resuspension and washes in PBS and cells were replated onto MS5 stroma for a further 5 week period, followed by methylcellulose addition.

### 9. Erythroid differentiation assay

K562 cells were subjected to erythroid differentiation by induction by DMSO (Fisher Scientific, Paisley, UK) over a period of 6 days, as described (2). 100 000 cells were seeded. At E0, 1.5% DMSO was added onto the culture medium (RPMI + 10% FBS + 1% P/S). RNA was collected at E0, E2, and E6 for subsequent qPCR analysis of erythroid gene markers *HBB, EPOR*, and *TAL1*.

### 10. RNA sequencing analysis

RNA sequencing was performed by Macrogen Europe BV (Amsterdam, Netherlands). A minimum of 1 μg (20 ng/μl) of high-quality total RNA (extracted using Monarch Total RNA Miniprep Kit, NEB) was supplied for sequencing. Macrogen Europe BV constructed libraries using Illumina Truseq Stranded Total RNA library preparation with Ribozero rRNA depletion, and performed sequencing was performedon a Novaseq 6000 platform, at 50M paired-end reads per sample. RNA-seq results in the form of fastq raw reads were analysed with the open source software package of the Tuxedo Suite. Tophat2 with Bowtie2 were used to map paired-end reads to the reference Homo sapiens genome build GRCh38 (5,6). GENCODE38 (7) was used as the reference human genome annotation. Aligned reads were filtered by quality using samtools (8) with a minimum selection threshold set at 30. Transcript assembly and quantification was achieved using Cufflinks (9). Differential expression between sample and control was performed by collapsing technical replicates for each condition and the use of the Cuffdiff tool (9). The differential expression was expressed in the form of log2 fold change between sample and control, and deemed statistically significant by a lower p value of 0.05 and false discovery rate (FDR) of 0.1.

### 11. Gene Ontology analysis

Gene ontology (GO) analysis was performed in ExpressAnalyst (available at www.expressanalyst.ca) using PANTHER Biological Process (BP) and Molecular Function (MF) repositories. GO terms and pathways were filtered by p value and false discovery rate (FDR) with a cut-off of ≤ 0.05 for meaningful association.

### 12. Statistical analysis and visualisation

Statistical significance was calculated by two-tailed Student’s t-test on available replicates (minimum n=3). Variance is represented in barcharts by error bars of ± standard deviation (SD). A p value ≤ 0.05 was considered statistically significant, unless otherwise stated. Statistical significance is symbolised as: p value ≤ 0.05 (*), 0.001 (**), 0.0001 (***), and 0.00001 (****). Calculations were performed in R studio version 4.1.0. Graphs were generated in Microsoft Excel 2013 and in R studio 4.1.0 using the libraries ggplot2 (v 3.3.5) and ggrepel (v 0.9.1). Multiplex Venn diagrams were generated on VennPainter (10).

### 13. Patient signature methods and data availability

Clinical phenotype and expression data (in counts units) were extracted from the Therapeutically Applicable Research to Generate Effective Treatments (TARGET, https://ocg.cancer.gov/programs/target) available in the GDC TARGET-AML cohort last accessed on 3^rd^ June 2022 from the University of California Santa Cruz (UCSC) Xena public repository (11). Microarray data in RMA units of expression for the TARGET AML cohort, including normal bone marrow data, was downloaded from the NCI TARGET Data portal. Microarray data in RMA units of expression for paediatric AML from Balgobind *et al*. (12) (under accession number GSE17855) was downloaded from GEO Accession Viewer. Data normalisation across arrays was achieved by Training Distribution Matching (TDM) using the R library TDM v 0.3 (13) and any remaining batch effects were removed by Limma removeBatchEffect. Differential expression analysis was performed using Limma for R (v 3.48.3). To construct the gene signatures, all differentially expressed genes for each comparison below a p value of 0.01 were included and intersected accordingly using Venn diagrams on VennPainter (10).

### 14. Gene Set Enrichment Analysis (GSEA)

Custom gene signatures were used as gene sets for GSEA analysis (Subramanian *et al*., 2005) on the GSEA software v4.2.3 on the RNA sequencing expression values in counts units of K562-t(7;12) against K562 control. GSEA was ran in 10 000 permutations on gene set using the weighted Signal2Noise metric. Enrichment plots and leading edge heatmaps were included as generated from the GSEA software.

